# Transcriptomic profiling of PBDE-exposed HepaRG cells unveils critical lncRNA-PCG pairs involved in intermediary metabolism

**DOI:** 10.1101/813337

**Authors:** Angela Zhang, Cindy Yanfei Li, Edward J. Kelly, Lianne Sheppard, Julia Yue Cui

## Abstract

Polybrominated diphenyl ethers (PBDEs) were formally used as flame-retardants and are chemically stable, lipophlic persistent organic pollutants which are known to bioaccumulate in humans. Although its toxicities are well characterized, little is known about the changes in transcriptional regulation caused by PBDE exposure. Long non-coding RNAs (lncRNAs) are increasingly recognized as key regulators of transcriptional and translational processes. It is hypothesized that lncRNAs can regulate nearby protein-coding genes (PCGs) and changes in the transcription of lncRNAs may act in *cis* to perturb gene expression of its neighboring PCGs. The goals of this study were to 1) characterize PCGs and lncRNAs that are differentially regulated from exposure to PBDEs; 2) identify PCG-lncRNA pairs through genome annotation and predictive binding tools; and 3) determine enriched canonical pathways caused by differentially expressed lncRNA-PCGs pairs. HepaRG cells, which are human-derived hepatic cells that accurately represent gene expression profiles of human liver tissue, were exposed to BDE-47 and BDE-99 at a dose of 25 μM for 24 hours. Differentially expressed lncRNA-PCG pairs were identified through DESeq2 and HOMER; significant canonical pathways were determined through Ingenuity Pathway Analysis (IPA). LncTar was used to predict the binding of 19 lncRNA-PCG pairs with known roles in drug-processing pathways. Genome annotation revealed that the majority of the differentially expressed lncRNAs map to PCG introns. PBDEs regulated overlapping pathways with PXR and CAR such as protein ubiqutination pathway and PPARα-RXRα activation but also regulate distinctive pathways involved in intermediary metabolism. BDE-47 uniquely regulated signaling by Rho Family GTPases and PBDE-99 uniquely regulates JAK/Stat signaling, bile acid biosynthesis, sirtuin signaling pathway, and autophagy. In conclusion, lncRNAs play essential roles in modifying important pathways involved in intermediary metabolism such as carbohydrate and lipid metabolism.

## INTRODUCTION

Polybrominated diphenyl ethers (PBDEs) are highly persistent organobromine compounds that were originally used as flame-retardants in a number of applications including textiles, plastics, and automobiles. There has been growing concern about the association between PBDE exposure and toxicity of the liver, thyroid, and nervous system. The sale of PBDEs was outlawed in 2003 in California and by the state of Washington in 2008 (http://www.leginfo.ca.gov/pub/03-04/bill/asm/ab_0301-0350/ab_302_bill_20030811_chaptered.html, https://www.doh.wa.gov/YouandYourFamily/HealthyHome/Contaminants/PBDEs). In 2004, the United States phased out the manufacture and import of the two most common PBDE mixtures: penta- and octa-BDE. Despite these measures, the United States Environmental Protective Agency (EPA) has reported increasing levels of PBDEs in humans and the environment (https://www.epa.gov/sites/production/files/2014-03/documents/ffrrofactsheet_contaminant_perchlorate_january2014_final_0.pdf). There are three potential sources for this increase: 1) the importation of products with PBDEs and 2) the degradation of PBDEs to more toxic and bioaccumulative congeners 3) the continued shedding of PBDEs into the environment from existing products. Despite its decreasing usage in commercial production, PBDEs will continue to continue to persist in the environment and contribute to adverse health concerns (Linares et al., 2015).

PBDEs bio-accumulate in the adrenal glands liver, kidneys, breast and adipose tissue through ingestion and inhalation (Linares et al., 2015). BDE-47 and BDE-99, in particular, were among the most dominant congeners found in both human tissue as well as indoor air and dust from US urban residences (Linares et al., 2015). Human breast milk specimens collected in North American over the last 15 years had total PBDE concentrations 20 times higher than samples collected in Europe or Asia (Zhang et al., 2017). Due to their small size, immature expression of xenobiotic detoxification genes, diet, and proximity to the ground, infants and toddlers are particularly vulnerable to potential developmental toxicity from PBDE exposure via ingestion and inhalation (Linares et al., 2015).

Although exposure to PBDEs can lead to neurotoxicity and the disruption of the endocrine system, the focus of this paper will be on impact on PBDEs on hepatotoxicity (Vuong et al., 2017 Eskenazi et al., 2013, Lilienthal et al., 2006). PBDEs have also been shown to play integral roles in oxidative stress and inflammation in the liver. Rats exposed to BDE-99 had increased superoxide mutase activity and oxidized glutathione levels, both of which are markers of oxidative stress (Albina et al., 2010). Furthermore, there is evidence in mouse models that BDE-47 increases liver weight and cytochrome P450 levels, which may induce a liver inflammatory response (Richardson et al., 2008). Hepatotoxicity has been suspected to be associated with PBDE-exposure, but there have been few studies that characterize the effect of PBDEs on human liver cells. Although many liver-derived cell lines exist, HepaRG cells are by far the most similar to human primary hepatocytes in terms of both gene expression levels and physical characteristics. Human primary hepatic cells rapidly lose their drug metabolizing functions shortly after suspension and poorly model xenobiotic metabolism in the liver (Andersson et al., 2012). Other commonly used human hepatic cell lines such as HuH7 and HepG2 lack accurate expression of drug processing genes. Originating from tumor in a female inflicted with hepatocarcinoma and hepatitis C, the HepaRG cell line is increasingly recognized as a superior method for accurate *in vitro* studies of the liver (Parent et al., 2004). This progenitor cell line is bipotent; HepaRG cells are able to differentiate into hepatocyte-like or biliary-like cells under certain defined conditions (Parent et al., 2004). A study in 2010 compared the gene expression of HepaRG and HepG2 cells against human primary hepatocytes using similarity matrices, principal components and hierarchical clustering methods. With regards to major drug processing genes such as cytochrome P450s, sulfotransferases, aldehyde dehydrogenases, and ATP-binding cassette transporters, the differences between expression in HepaRG and HepG2 cells were much larger than the differences found between HepaRG cells and primary human hepatocytes (Hart et al., 2010). From these conclusions, the HepaRG cell line more accurately represents human hepatic cells and should be considered for *in vitro* studies concerning PBDE exposure.

PBDEs activate the xenobiotic-sensing nuclear receptors (NR) pregnane X receptor (PXR) and constitutive androstane receptor (CAR) (Pacyniak et al., 2007; Sueyoshi et al., 2014). PXR and CAR are highly expressed in liver and transcriptionally regulate the expression of target genes by recruiting co-activators or co-repressors (Chai et al., 2013). As a ligand-dependent transcription factor, PXR is activated by a wide range of chemicals including glucocorticoids, antibiotics, and antifungals. Importantly, PXR is the principal transcriptional trans-activator of cytochrome P450 3A (CYP3A), an important class of oxidizing enzymes for metabolizing drugs and other xenobiotics. Similar to PXR, CAR is a transcriptional regulator of the cytochrome P450 2B6 (CYP2B6) enzyme. Although CYP2B6 has a minor role in drug metabolism compared to CYP3A4 (a major CYP3A isoform for drug metabolism), there may be crosstalk between PXR and CAR in regulating the expression of P450 genes (Willson and Kliewer, 2002). Rifampicin (RIF) is a unique activator of PXR and 6-(4-chlorophenyl)imidazo[2,1-b][1,3]thiazole-5-carbaldehydeO-(3,4-dichlorobenzyl)oxime (CITCO) is a unique activator of CAR. As a result, these compounds can be used to delineate the effect PBDEs have on PXR-CAR pathways (Timsit and Negishi, 2007).

While much attention has been focused on the impact of protein-coding genes (PCGs) in xenobiotic metabolism of the liver, there is growing evidence that long non-coding RNAs (lncRNAs) play important roles in the regulation of transcriptional and translation processes of nearby protein coding genes. LncRNAs, which are over 200 nucleotides in length, are transcribed by RNA polymerase II from the genome and are highly tissue-specific and developmentally regulated (Dempsey and Cui, 2017). Although they are not translated into protein, lncRNAs can share similar characteristics and mechanisms found in messenger RNA (mRNA) such as alternative splicing and polyadenylation. LncRNAs exert their function as gene regulators by serving as signals, decoys, and scaffolds through a partnership with RNA binding proteins in ribonucleoprotein complexes (He et al., 2019).

Several lncRNAs have been identified to play regulatory roles following exposure to toxicants. For example, high levels of phthalate metabolites have been associated with decreased levels of methylation at the *H19* loci. H19, a lncRNA, plays important regulatory roles in fetal and placental growth during development and is typically silenced postnatally (LaRocca et al., 2014). H19 can be re-activated in adulthood during tissue-regeneration; however, increased levels of H19 have been associated with tumor development in many cancers, including hepatocellular carcinoma. In a mouse study, levels of H19 RNA were increased following hypoxia and was followed by the nearly compete attenuation of p57^kip2^, a KIP family cyclin-dependent kinase (Cdk) inhibitor that holds control over the cell cycle (Matouk et al., 2007). A large number of lncRNAs were dysregulated in liver of a mouse model following exposure to PBDEs, suggesting a possible link between the regulation of lncRNAs and PBDE-mediated toxicity (Li and Cui, 2018). Little is known regarding the effect of PBDE exposure on lncRNAs in human liver, which is a major organ for xenobiotic metabolism and nutrient homeostasis.

Taken together, the goals of this study are 1) to characterize what PCGs and lncRNAs are differentially regulated following PBDE exposure, 2) to identify PCG-lncRNA pairs through gene annotation and predictive binding tools, and 3) to determine enriched canonical pathways caused by differentially expressed lncRNA-PCG pairs.

## METHODS

### HepaRG Cell Culture

The undifferentiated HepaRG cells were provided by Dr. Edward J. Kelly (School of Pharmacy, University of Washington, Seattle, WA), which were initially purchased from Biopredic International (Rennes, France). The HepaRG cells were seeded at 2.6 × 10^4^ cells/cm^2^ in six-well plates and grown in William’s medium E supplemented with growth medium penicillin, and 100 μg/ml streptomycin. After 2 weeks, the cells were shifted to the same medium supplemented with differentiation medium supplement (Catalog # ADD 721, Triangle Research Labs, NC), 100 IU/ml penicillin, and 100 μg/ml streptomycin to differentiate the cells into a hepatocyte-like morphology. The cells were cultured in differentiation medium for another 2 weeks. The medium was renewed every 2 to 3 days. Prior to the treatment, the cell medium was changed to induction medium (Williams’ medium E with induction supplement (HPRG740, Life Technologies, Carlsbad, CA) for 24 h. The fully differentiated HepaRG cells were then treated with vehicle control (0.1% DMSO), CITCO (1 µM), RIF (10 µM), BDE-47 (25 µM), or BDE-99 (25 µM) in triplicates for 24 hours. Total RNA from these treated cells was isolated using RNA-Bee reagent (1 ml/well) per manufacturers’ instructions. During this experiment, we observed no cytotoxicity of BDE-47 or BDE-99 on HepaRG cells based on the MTT viability assay (data not shown).

### RNA-sequencing analysis

The workflow for the analysis is outlined in Figure 1. FASTQ files with paired-end sequence reads were mapped to the human genome (hg19) using HISAT (Hierarchical Indexing for Spliced Alignment of Transcripts). The resulting SAM (sequence alignment/map) files were converted to its binary form (BAM: binary alignment/map) and sorted using SAMtools (verison 1.2). PCG and lncRNA transcript abundances were estimated with featureCounts (part of the Subread package, version 1.5.3) using the UCSC hg19 and NONCODE 2016 database as reference annotation. Transcript abundances were expressed as integer counts and transcripts with less than 3 reads per sample were excluded from further analysis. Differential analysis with DESeq2 (Version 1.16.1) (Love et al., 2014) was performed for each treatment against the control. Transcripts were considered differentially regulated by treatment if their Benjamini-Hochberg adjusted false discovery rate (FDR-BH) was less than 0.05. A differentially expressed genes was defined as being significantly different from control (FDR <0.05) in at least one of the chemical exposure groups (BDE-47, BDE-99, CITCO, and RIF) and having at least 3 reads per sample. A stably expressed gene was defined as having >3 reads per sample, and its expression not significantly different from control in any of the chemical exposure groups. Venn diagrams comparing the set of differentially expressed transcripts in each treatment were generated for the lncRNAs and PCGs using the ‘gplots’ package (version 3.0.1) in R.

**Figure 1:**
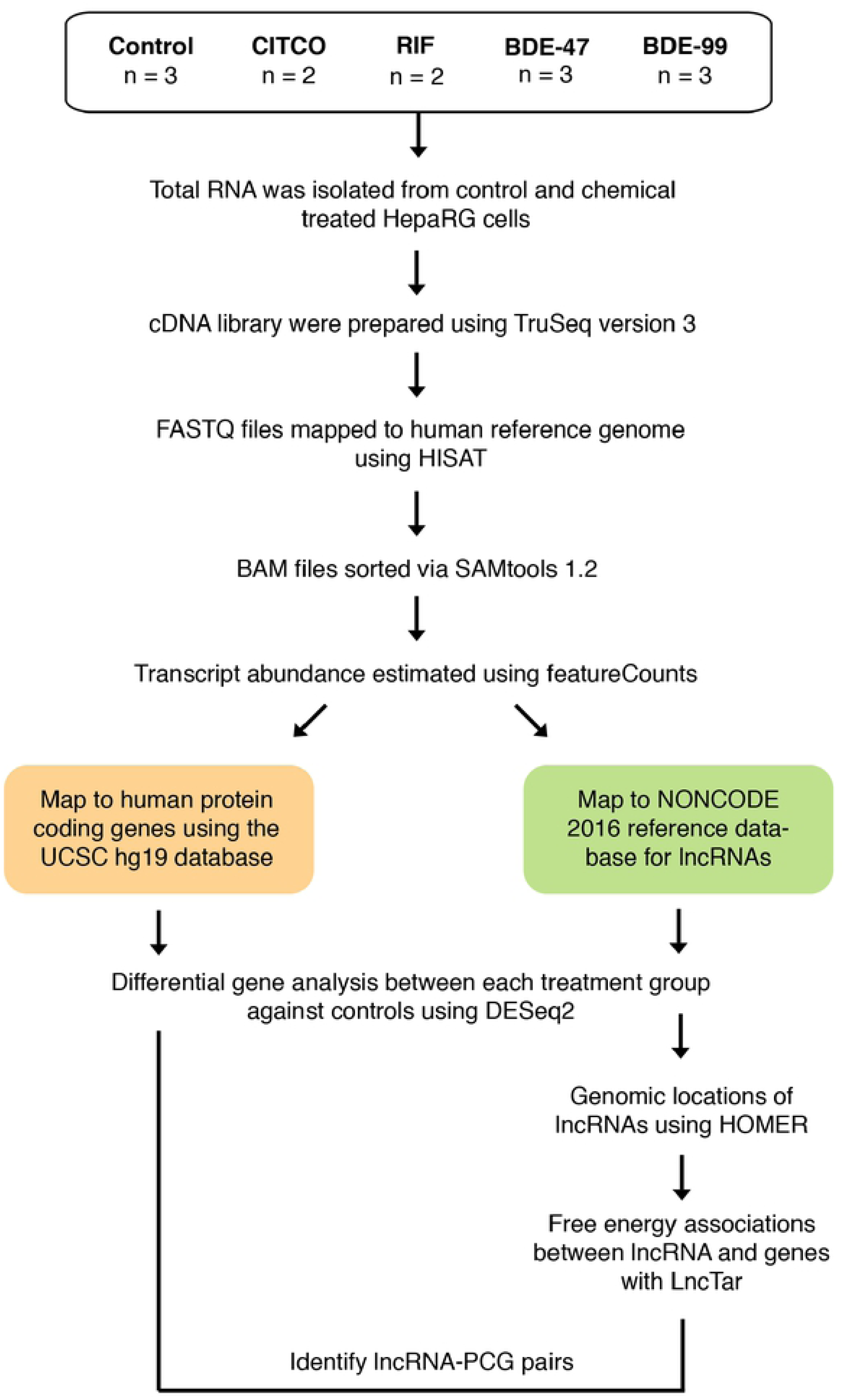
Workflow.

### Genomic annotation of lncRNAs and identification of proximal lncRNA-PCG pairs

The command line tool, annotatePeaks, which is part of the HOMER (Hypergeometric Optimization of Motif EnRichment, version 4.9) suite was used to identify lncRNAs proximal to PCGs using UCSC hg19 as a reference database. LncRNAs were considered proximal to a PCG if they were in the transcription start site (TSS, defined from −1 kb to +100 bp), transcription termination site (TTS, −100 bp to +1 kb), 5’ untranslated region exon, 3’ UTR Exon, or in an intron. LncRNA-PCG pairs within treatment were identified if annotatePeaks assigned a differentially expressed lncRNA to a differentially expressed PCG.

### Pathway analysis of differentially expressed lncRNA-PCG pairs

Gene enrichment analysis by treatment was performed on the differentially expressed PCGs that were proximal to differentially expressed lncRNAs through Ingenuity Pathway Analysis (IPA). The significant canonical pathways from each treatment were pooled together and ordered by level of significance. Ten canonical pathways were chosen by these criteria and were all broadly under the categories of Nuclear Receptors, Immune Response or Protein Ubiquitination. Gene expression heatmaps were plotted for all four treatments in each canonical pathway using the ‘made4’ package (version 1.50.0) in R. Venn diagrams comparing the set of genes in each treatment by pathway was plotted using the ‘gplots’ package (version 3.0.1) in R.

### LncRNA-PCG binding prediction with LncTar

Human lncRNA sequences were retrieved from the NONCODE 2016 database (http://www.noncode.org/download.php) and human protein-coding transcript sequences were retrieved from Ensembl Biomart. LncTar Version 1.0 was used to generate a tab-delimited file of lncRNA and PCG pairs based on predicted free-energy associations between nascent transcripts. A threshold of −0.08 normalized free energy (ndG: a novel parameter set by lncTar) was set because it is the lowest suggested threshold to detect all possible lncRNA-mRNA interactions.

## RESULTS

The goal of this study was to identify functional interactions between PCGs and lncRNAs through RNA-Seq in HepaRG cells following PBDE exposure.

### Regulation of PCGs and lncRNAs following exposure to PBDEs

In order to identify functional interactions between PCGs and lncRNAs RNA-Seq was performed on HepaRG cells exposed to BDE-47 and BDE-99. We also exposed HepaRG cells to CITCO (a CAR agonist) and RIF (a PXR agonist) in order to identify any overlapping PXR/CAR pathways with PBDEs. Table 1 shows that ∼80-125 million reads (95-99% of total reads) were mapped to the human reference genome (NCBI GRCh37/hg19). Based on these filtering criteria, 19.3% of the total annotated genes were not expressed in any exposure groups, 47.7% were stably expressed, and 33.1% were differentially expressed by chemical exposure (Figure 2A).

**Figure 2:**
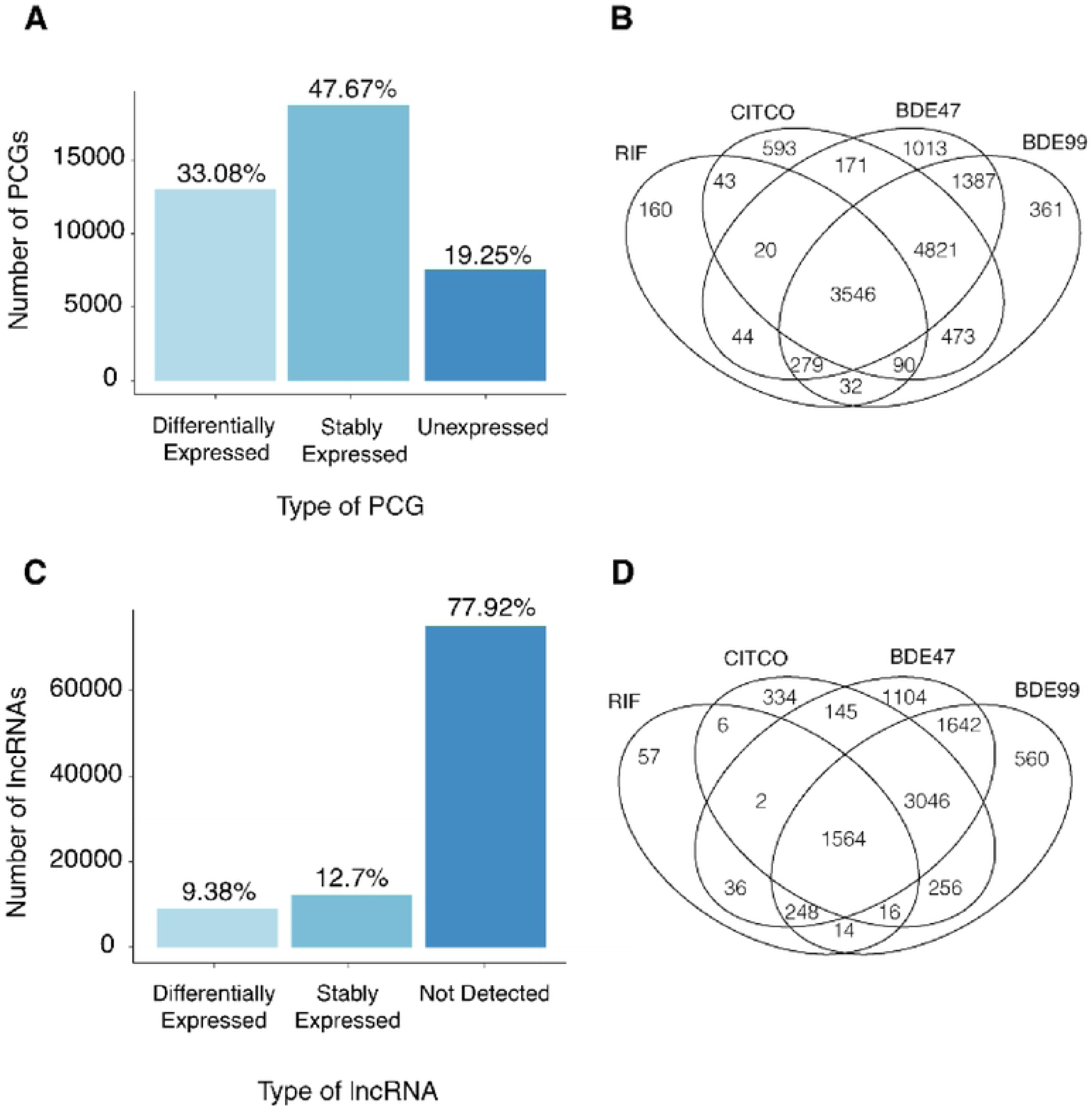
Regulation of protein coding genes (PCGs) and long noncoding RNA (lncRNAs) in HepaRG cells when exposed to rifampin (RIF), constitutive androstane receptor agonist (CITCO), 8DE47 and BDE99. (A) Bar plot depicting the number of PCGs that were differentially expressed in at least one treatment group, stably expressed in all treatment groups, and unexpressed in all treatment groups. (B) Four-way Venn Diagram displaying shared and unique differentially expressed PCGs in all treatment groups. (C) Bar plot depicting the number of lncRNAs that were differentially expressed in at least one treatment group, stably expressed in all treatment groups, and unexpressed in all treatment groups. (D) Four-way Venn Diagram displaying shared and unique differentially expressed lncRNAs in all treatment groups.

A four-way Venn diagram (Figure 2B) was generated from the group of differentially expressed PCGs to identify unique and common pathways among all four chemical exposures. There were 3546 commonly regulated PCGs among the BDE-47, BDE-99, CITCO, and RIF exposure groups, whereas a larger group of 4821 genes was commonly regulated among CITCO and the PBDE groups. In contrast, there was less overlapping between RIF exposed group and the PBDE exposed groups. BDE-47 had 1013 uniquely regulated genes and BDE-99 had 361 uniquely regulated genes. In summary, there is a large overlap of PCGs in all four chemical exposure groups, suggesting that PBDEs and PXR/CAR activation affect similar pathways. The even larger overlap of PCGs between CITCO and PBDEs exposed groups indicate that PBDE exposure may activate CAR more than PXR signaling.

Regarding lncRNAs, a majority (77.9%) were not expressed in any exposure groups, 12.7% were stably expressed, and 9.38% were differentially expressed in at least one chemical exposure group (Figure 2C). A 4-way Venn diagram (Figure 2D) for the differentially expressed lncRNAs showed similar regulatory patterns as compared to the PCGs: 1564 lncRNAs were differentially expressed in by all four chemical exposures. A larger group of 3046 lncRNAs were commonly regulated among the PBDEs and CITCO exposure groups. A total of 1104 lncRNAs were uniquely regulated by BDE-47 whereas only 560 lncRNAs were uniquely regulated by BDE-99.

### Genomic annotation of differentially regulated lncRNAs relative to PCGs

All differentially expressed lncRNAs were annotated using the annotatePeaks function from the HOMER (Hypergeometric Optimization of Motif EnRichment) suite. A vast majority (72.1%) of lncRNAs were mapped proximal to the introns of PCGs followed by the exons (7.77%), the 3’-untranslated regions (3’-UTRs) (6.75%), the transcription start sites (TSS) (5.95%), promoter (5.15%) and the 5’-UTR (0.49%) of PCGs (Figure 3).

**Figure 3:**
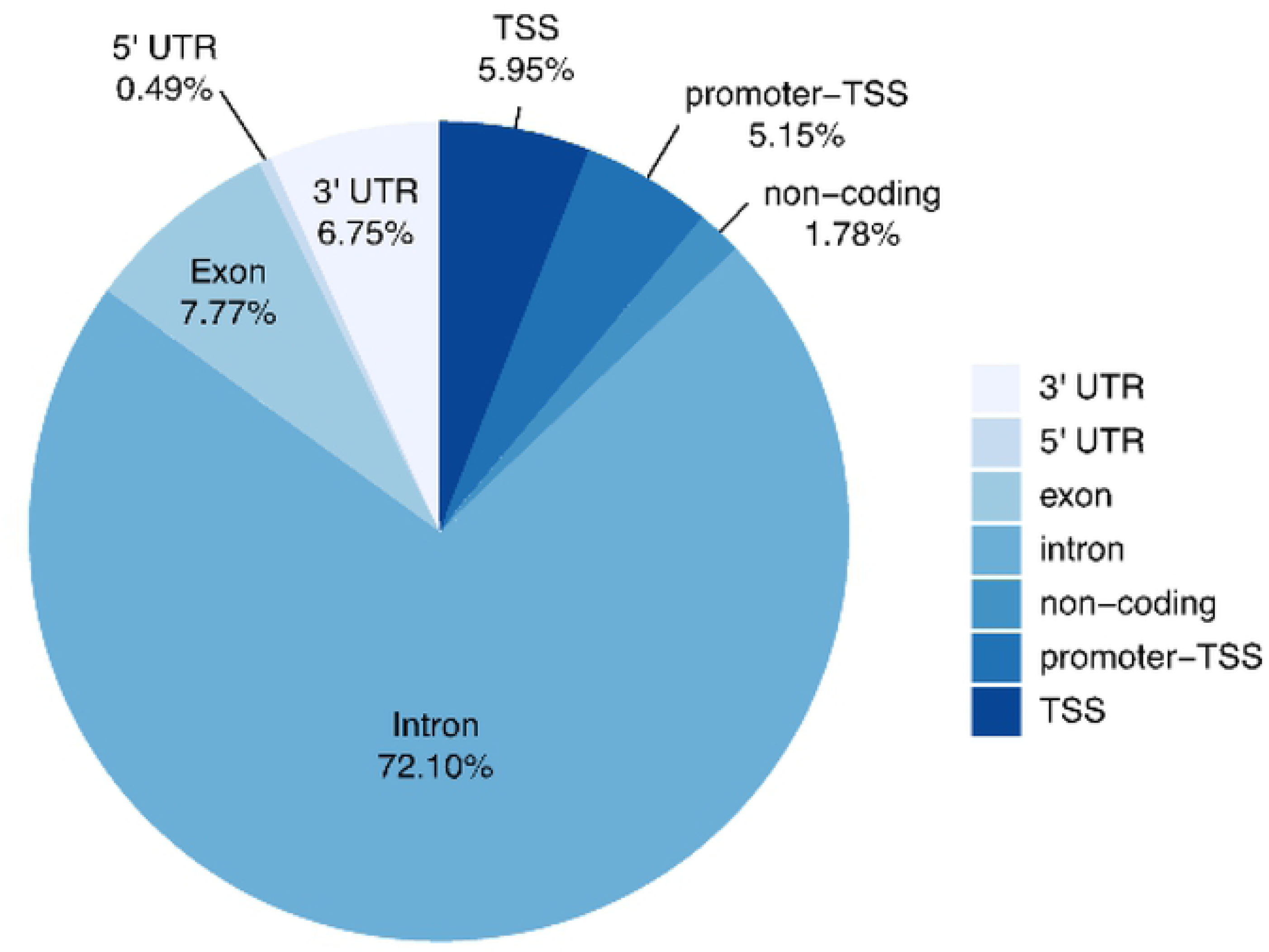
Genome annotation of lncRNAs. Pie chart showing the proportion of lncRNAs that were mapped proximal to either a 3’UTR, 5’UTR, exon, intron, non­ coding region, promoter TSS, or TSS.

### Enriched canonical pathways following exposure to PBDEs, CITCO and rifampicin

PCG-lncRNA pairs were determined through two criteria: 1) both PCG and lncRNA were differentially expressed by the same exposure group and 2) the differentially expressed lncRNA mapped to the differentially expressed PCG.

#### Protein Ubiquitination Pathway

The protein ubiquitination pathway was significantly disrupted in all four exposure groups. This pathway is responsible for the degradation of an ubiquitin-protein conjugate by proteasomes. There were 15 PCGs in this pathway that were commonly regulated by CITCO, RIF, BDE-47, and BDE-99 (Figure 4A). Several members of the Ubiquitin Specific Peptidase family (USP24, USP7, USP37, USP40, USP25) were up-regulated by all four chemicals. These proteins belong to a large family of cysteine proteases which deubiquitinate and reverse the degradation of proteins. In particular, USP7 deubiquitinates p53 and WASH (part of the Wiskott–Aldrich Syndrome protein family), suggesting its role in disrupting tumor suppression pathways (Zhou et al., 2018). Several PCGs associated with heat-shock proteins (HSPs) such as DNAJC13, Sacsin Molecular Chaperone (SACS), HSPA4, and HSPA4L were up-regulated. The expression of Ubiquitin Protein Ligase E3 Component N-Recognin 1 (UBR1), Cullin 2 (CUL2), and mouse double minute 2 homolog (MDM2), genes encode components for the E3 ubiquitin ligase, were also up-regulated by all four treatment groups. MDM2, in particular, can promote tumor formation by targeting p53, a tumor suppressor, for degradation (Zhou et al., 2018). General vesicular transport factor p115 (USO1), a gene that encodes for a peripheral membrane protein that recycles between the cytosol and Golgi apparatus during interphase, was up-regulated in all four treatments as well. Out of the 15 commonly regulated genes, only two, namely the DnaJ Heat Shock Protein Family (Hsp40) Member C4 (DNAJC4) and the 26S proteasome non-ATPase regulatory subunit 8 (PSMD8), were down-regulated. DNAJC4 is related to unfolded protein binding, whereas down-regulation of PSMD8 is known to lead to the accumulation of damaged or misfolded protein species (Kleiger and Mayor, 2014) (Figure 4B-E).

**Figure 4:**
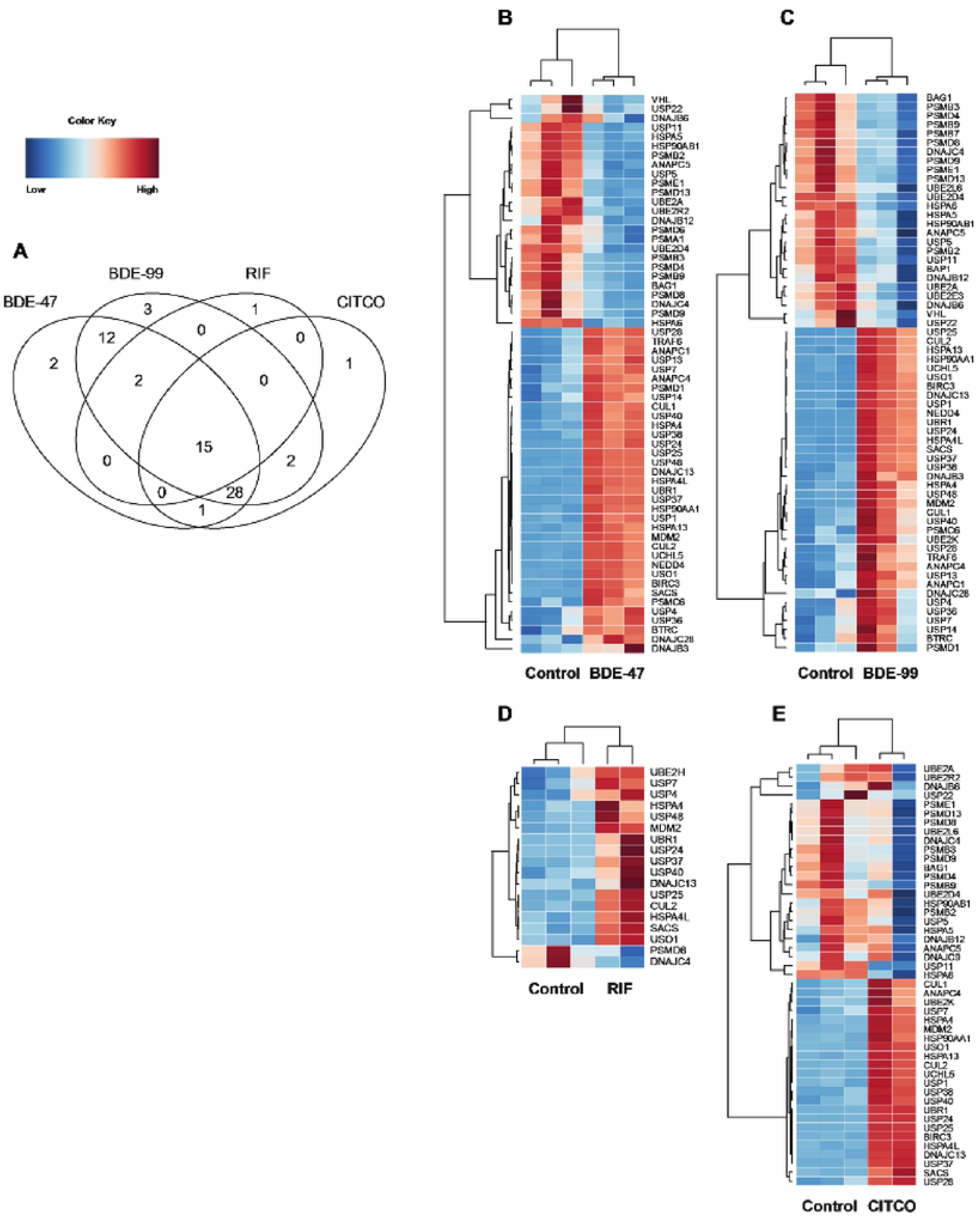
Differentially expressed lncRNA-PCGs pairs associated with the Protein Ubiquitination pathway: (A-B) Heatmaps depicting gene expression patterns of differentially expressed genes associated with this pathway. Blue indicates down-regulation while red indicates up-regulation. (E) Four-way Venn Diagram displaying the common and unique genes differentially expressed in all four pathways.

There were several genes that were uniquely regulated in each exposure group. 26S proteasome non-ATPase regulatory subunit 6 (PSMD6) and Proteasome Subunit Alpha 1 (PSMA1), two genes involved in the degradation of ubiquinated proteins, were down-regulated only by BDE-47 (Figure 4B). PSMD6 encodes a member of the protease subunit S10 family while PSMA1 encodes a component of the 20S core proteasome complex and plays a key role in removing misfolded or damaged proteins in order to maintain protein homeostasis (Kleiger and Mayor, 2014). Ubiquitin Conjugating Enzyme E2 E3 (UBE2E3), Proteasome Subunit Beta 7 (PSMB7), and BRCA1 Associated Protein 1 (BAP1) were down-regulated only by BDE-99 (Figure 4C). UBE2E3 accepts ubiquitin from the E1 complex and attaches to proteins targeted for degradation. BAP1 belongs to a subfamily of deubiquitinating enzymes that removes ubiquitin from proteins and is a tumor suppressor of BRCA1. Ubiquitin Conjugating Enzyme E2 H (UBE2H), part of a family of ubiquitin-conjugating enzymes, was up-regulated in RIF (Figure 4D) and DnaJ Heat Shock Protein Family (Hsp40) Member C9 (DNAJC9) was down-regulated in CITCO (Figure 4E).

#### PPARα-RXRα Pathway

Peroxisome proliferator-activated receptor alpha (PPARα) belongs to a subfamily of transcription factors and plays a major role in lipid metabolism. It regulates mitochondrial and peroxisomal fatty acid oxidation during prolonged deprivation of food (Kersten et al., 1999). Following exposure to all four treatments, cytochrome P450 family 2 subfamily C member 8 (CYP2C8) and cytochrome P450 family 2 subfamily C member 9 (CYP2C9) were up-regulated compared to control. CYP2C9 makes up 18% of all cytochrome P450 proteins in liver; it is considered to be one of the most important drug-metabolizing enzymes and plays a critical role in metabolizing around 15% of all clinically important drugs (Dai et al., 2015). CYP2C8 is responsible for metabolizing many xenobiotics such as mephenytoin, benzo(a)pyrene, and taxol and also converts arachidonic acid to epoxyeicosatrienoic acid in the human liver and kidney (Dai et al., 2001) (Figure 5A).

**Figure 5:**
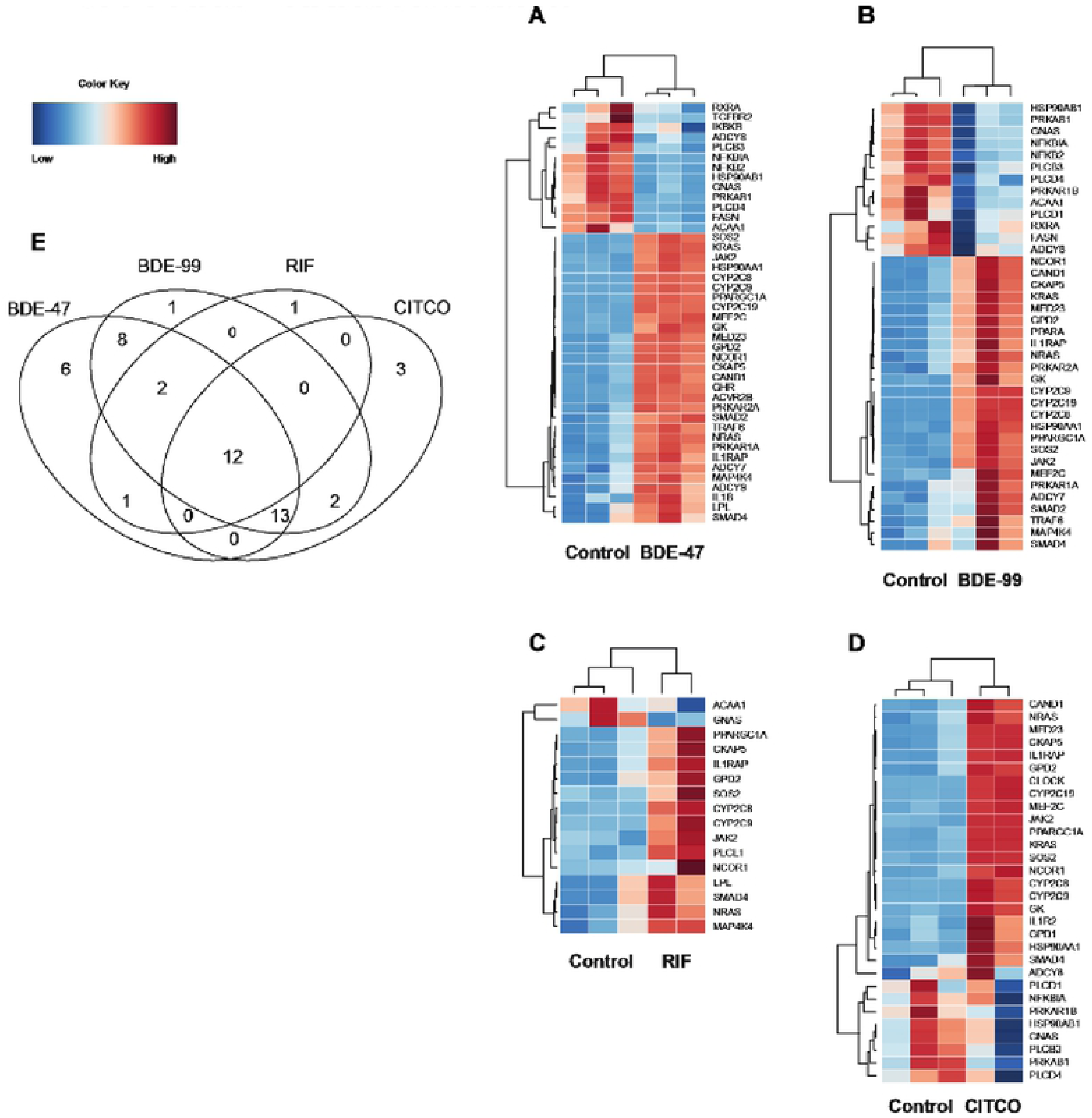
Differentially expressed lncRNA-PCGs pairs associated with the PPARα-RXRα pathway: (A-D) Heatmaps depicting gene expression patterns of differentially expressed genes associated with the PPARα -RXRα pathway. Blue indicates down-regulation while red indicates up-regulation. (E) Four-way Venn Diagram displaying the common and unique genes differentially expressed in all four pathways.

SMAD family member 4 (SMAD4), nuclear receptor corepressor 1 (NCOR1) and PPARG coactivator 1 alpha (PPARGC1A) were also up-regulated in BDE-47, BDE-99, RIF, and CITCO groups (Figure 5A). SMAD4 belong to a family of transcription factor proteins and mediate TCF-β signal transduction. These interactions are critical for regulating important processes such as embryo development, tissue homeostasis, and immune regulation (Massague, 2012). NCOR1 mediates transcriptional repression and belongs to a complex that promotes the formation of chromatin structures that downregulate gene expression (Muller et al., 2017). SMAD4 and NCOR1 are two cancer driver genes associated with breast cancer, carcinoma, and brain cancer (Ledgerwood et al., 2016; Pereira et al., 2016; Tsigelny et al., 2016). PPARGC1A is a transcriptional coactivator that regulates genes involved in energy metabolism. It links external physiological stimuli with mitochondrial biogenesis and plays an important contribution in muscle fiber type determination. NCOR1 and PPARGC1A oppose each other: PPARGC1A enhances PPAR gamma activity while NCOR1 represses it (He et al., 2015).

The PPARα-RXRα pathway play important roles in cell bioenergetics, inflammatory responses, and cellular structure in the liver (Liss and Finck, 2017; Patsouris et al., 2004; Pawlak et al., 2015). Several genes found in these pathways were up-regulated in all four treatment groups. Glycerol-3-phosphate dehydrogenase 2 (GPD2) is involved in glycerol production through the glycerol kinase pathway. Located in the mitochondria, GPD2 plays a pivotal role in cell bioenergetics and functions in the intersection of glycolysis, oxidative phosphorylation, and fatty acid metabolism (Mracek et al., 2013). Interleukin 1 Receptor Accessory Protein (IL1RAP) induces the synthesis of acute phase proteins. During infection, tissue damage, or stress, IL1RAP activates interleukin 1-responsive genes, which then form a complex in the cell membrane (Wesche et al., 1997). Cytoskeleton associated protein 5 (CKAP5) belongs to the transporter-opsin-G protein-coupled receptor (TOG) family and plays an important role in spindle formation by protecting kinetochore microtubules from depolymerization (Lu et al., 2017).

Only one gene was down-regulated following exposure to BDE-47, BDE-99, RIF, and CITCO. heterotrimeric G-protein alpha subunit complex locus (GNAS) has highly complex imprinted expression pattern encodes for the stimulatory G-protein alpha subunit. GNAS is a signal transducer and transmits hormonal and growth factor to downstream proteins (Zauber et al., 2016) (Figure 5A).

Six genes were uniquely regulated by exposure to BDE-47 (Figure 5B). Two genes involved in tumorigenesis, transforming growth factor beta receptor 2 (TGFBR2), and inhibitor or nuclear factor kappa B kinase subunit beta (IKBKB) were down-regulated compared to the control. TGFBR2 is a transmembrane protein that forms a heterodimeric complex with TGF-beta receptor type 1. This complex phosphorylates proteins that regulate the transcription of genes related to cell proliferation, cell cycle arrest, and apoptosis (Wei et al., 2015). IKBKB activates NF-kB and promotes tumorigenesis through the inhibition of forkhead transcription factor FOXO3 (Hu et al., 2004). Interleukin 1 Beta (IL1B), an important mediator of the inflammatory response, was up-regulated. It is hypothesized that inhibition of NF-kB pathway intermediates such as IKBKB could result in a reduction of pro-inflammatory genes like IL1B (Douglass et al., 2017).

Although TGFBR2 was down-regulated by BDE-47, activin receptor type-2B (AVCR2B), another member of the transforming growth factor-beta superfamily, was up-regulated. AVCR2B activates activin, which in turn regulates follicle-stimulating hormone production by pituitary cells (Attisano et al., 1996). Growth hormone receptor (GHR), a member of the type I cytokine receptor family, was also up-regulated. Binding of growth hormone and GHR activates the intra- and intercellular signal transduction pathway and leads to growth. Interleukin 1 Beta (IL1B) was up-regulated along with GHR, and there is evidence that pro-inflammatory cytokines such as IL-1β are involved in hepatic growth hormone resistance during inflammation (Zhao et al., 2014).

Peroxisome proliferator activated receptor alpha (PPARA) was up-regulated only in BDE-99 treated samples (Figure 5C). Peroxisome proliferators include hypolipidemic drugs, plasticizers, and herbicides. These peroxisome proliferators are mediated via specific receptors such as PPARA, which in turn affects target genes involved in cell proliferation, cell differentiation and inflammation responses (Tyagi et al., 2011). Phospholipase C Like 1, inactive (PLCL1) was up-regulated in response to RIF exposure (Figure 5D). PLCL1 is involved in the inhibition of exocytosis through its interactions with syntaxin 1 and SNAP-25 (Zhang et al., 2013b).

Three genes were uniquely up-regulated in response to CITCO (Figure 5E). Glycerol-3-Phosphate Dehydrogenase 1 (GPD1) plays an essential role in carbohydrate and lipid metabolism and converts dihydroxyacetone phosphate (DHAP) and reduced nicotine adenine dinucleotide (NADH) to glycerol-3-phosphate and NAD+ (Ou et al., 2006). Interleukin 1 Receptor Type 2 (IL1R2) acts as a decoy receptor that inhibits the activity of interleukin alpha (IL-1α) and interleukin beta (IL-1β). Clock circadian regulator (CLOCK) plays a central role in the regulation of circadian rhythms. It is hypothesized that melatonin affects inflammatory cytokines such as a IL1R2 which in turn may affect a mechanism involving CLOCK (Kowalewska et al., 2017).

### PBDEs uniquely affect pathways related to carbohydrate metabolism

The GDP-L-fucose biosynthesis I (from GDP-D-mannose) pathway was uniquely down-regulated following exposure to both BDE-99 and BDE-47 (Figure 6A-B). GDP-L-fucose is the activated nucleotide sugar form of L-fucose and is integral to the structure of many polysaccharides and glycoproteins such as N-linked glycans (Bonin et al., 1997). GDP-mannose 4,6-dehydratase (GMDS) and GDP-L fucose synthetase (TSTA3) were down-regulated by PBDE exposure. GMDS is involved in the first of this two-step process and converts GDP-D-mannose to GDP-4-dehydro-6-deoxy-D-mannose through an epimerase reaction. TSTA3 completes the pathway and synthesizes GDP-L-fucose from GDP-4-dehydro-6-deoxy-D-mannose via a reductase reaction (Tonetti et al., 1996).

**Figure 6:**
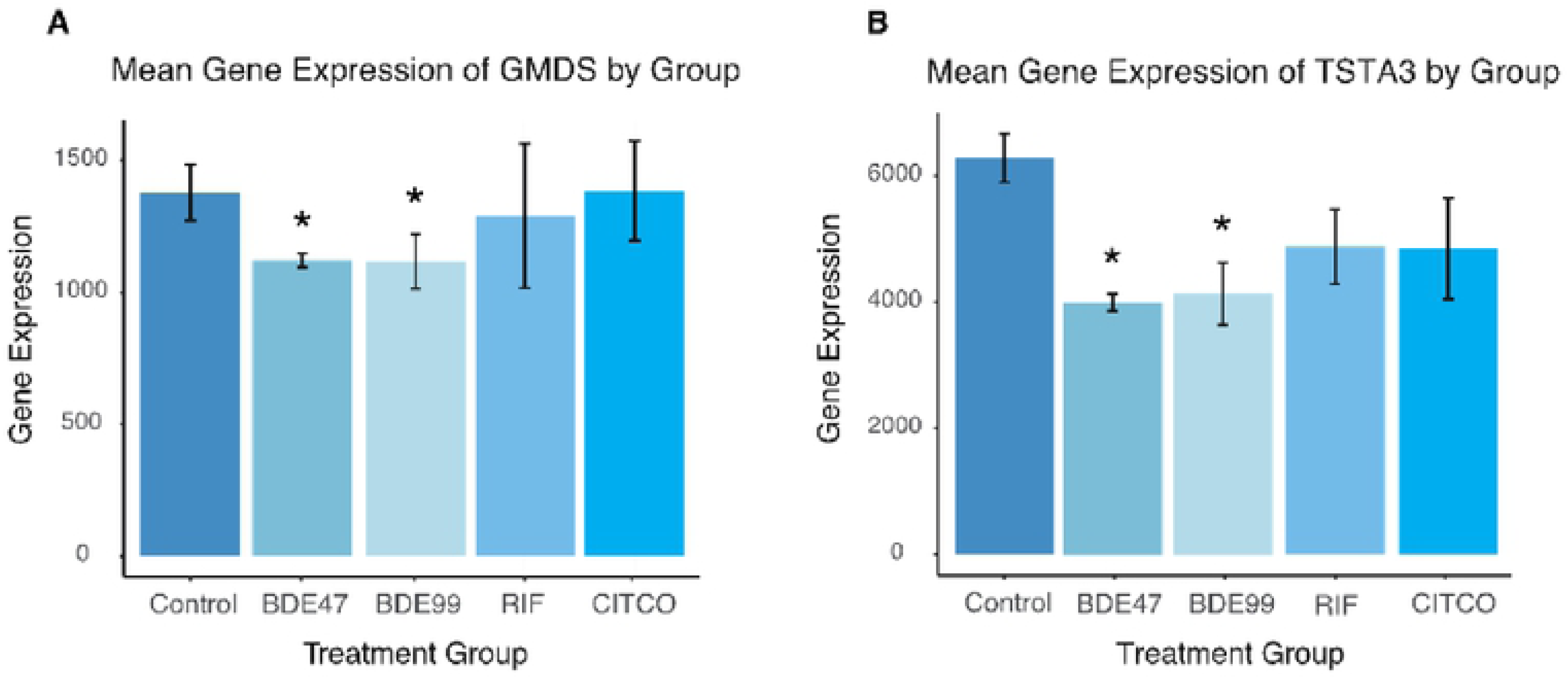
Differentially expressed lncRNA-PCGs pairs associated with the GDP-L­ fucose Biosynthesis I (from GDP-D-mannose) pathway: (A-B) Barplots depicting gene expression patterns of differentially expressed genes associated with the GDP-L-fucose Biosynthesis I pathway. Errors bars represent mean ± SE.

### Enriched canonical pathways involving Rho family of GTPases following exposure to BDE-47

Rho family of GTPases contains a superfamily of proteins that act as molecular switches for signal transduction pathways that drive many dynamic aspects of cell behavior. Ras homolog gene family, member A (RHOA), ras-related C3 botulinum toxin substrate 1 (RAC1), and cell division control protein 42 homolog (CDC42) are perhaps the most well-studied proteins in this superfamily and convert between inactive GDP-bound and active GTP-bound conformational states (Hall, 2012). Following exposure to BDE-47, there was a decrease in myosin light chain genes (MYL5, MYL6, MYL12A) and an increase in cadherin genes (CDH7, CDH19, CDH10, CDH12). Cadherins, which aid in the formation of cell-cell adherens junctions, dramatically decreases RHOA activity and stimulates CDC42 and RAC1 activity (Noren et al., 2001). Furthermore, literature suggests that a decrease of Rho family GTPs causes a decrease in the expression of MYL genes (Salhia et al., 2008). BDE-47 down-regulated the expression of Ras homology family member T2 (RHOT2) and Ras homolog family member D (RHOD); RHOT2 and RHOD interact with protein kinases and may be targets for activated GTPases (Ridley, 2006). Furthermore, the expression Rho guanine nucleotide exchange factors and Rho GTPase activating proteins (ARHGEF5, ARHGEF19, ARHGAP9), which stimulate the GTP-GDP exchange reaction, were down-regulated. Although many genes involved in the Rho family of GTPases signaling pathway were down-regulated, ARHGEF11, ARHGAP12, and Rho-associated protein kinase 2 (ROCK2) were up-regulated (Figure 7A-B).

**Figure 7:**
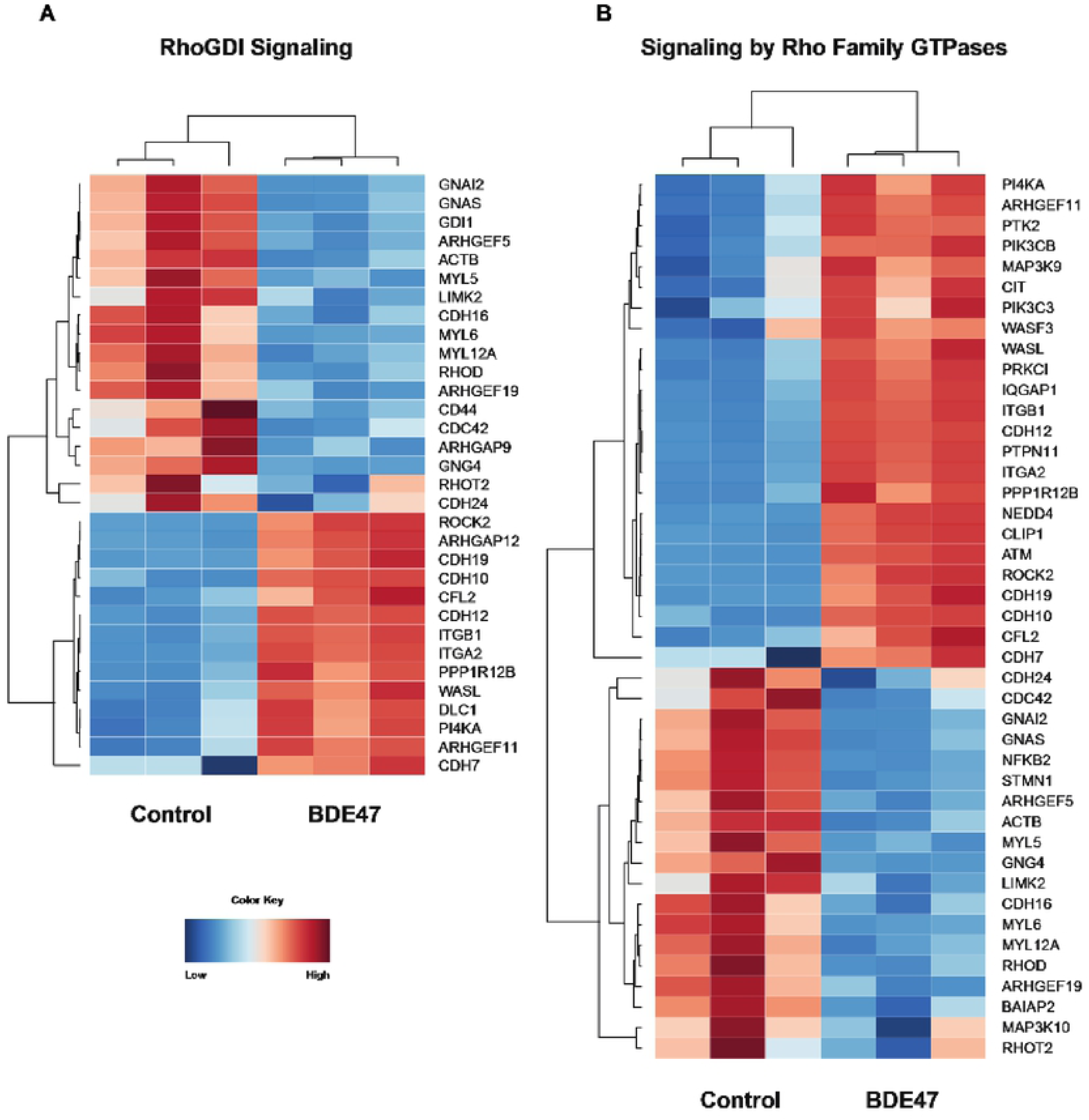
Effect of BDE47 on Rho Family of GTPases: (A-B) Heatmaps depicting gene expression patterns of differentially expressed genes associated with the RhoBDI Signaling pathway (A) and Signaling by Rho Family GTPases (B). Blue indicates down­ regulation while red indicates up-regulation.

### Enriched canonical pathways following exposure to BDE-99

#### Bile Acid Synthesis Pathway

As shown in Figure 8A, following exposure to BDE-99, bile acid-CoA:amino acid N-acyltransferase (BAAT), cytochrome P450 family 3 subfamily A member 4 (CYP3A4), and aldo-keto reductase family 1 member C4 (AKR1C4) were up-regulated while cytochrome P450 family 27 subfamily A member 1 (CYP27A1) and hydroxy-delta-5-steroid dehydrogenase, 3 Beta- And Steroid delta-isomerase 7 (HSD3B7) was down-regulated. BAAT is a liver enzyme that catalyzes the transfer of C24 bile acids to glycine or taurine (Styles et al., 2016), and AKR1C4 is involved in downstream of bile acid synthesis in hepatocytes (Rizner and Penning, 2014). CYP3A4 is a prominent drug-processing gene involved in the metabolism of a variety of commonly used drugs and is thought to contribute to bile acid metabolism during liver diseases (Zanger and Schwab, 2013). CYP27A1 is also a member of the cytochrome P450 superfamily of enzymes and plays an important role in cholesterol homeostasis (Zanger and Schwab, 2013). Whereas HSD3B7 is involved in the initial stages of bile acid synthesis and is encodes a membrane-associated endoplasmic reticulum protein (Schwarz et al., 2000). In summary, exposure to BDE-99 caused a down-regulation in genes associated with the initial stages of bile acid synthesis and an up-regulation of genes associated with the later stages of bile acid synthesis.

**Figure 8:**
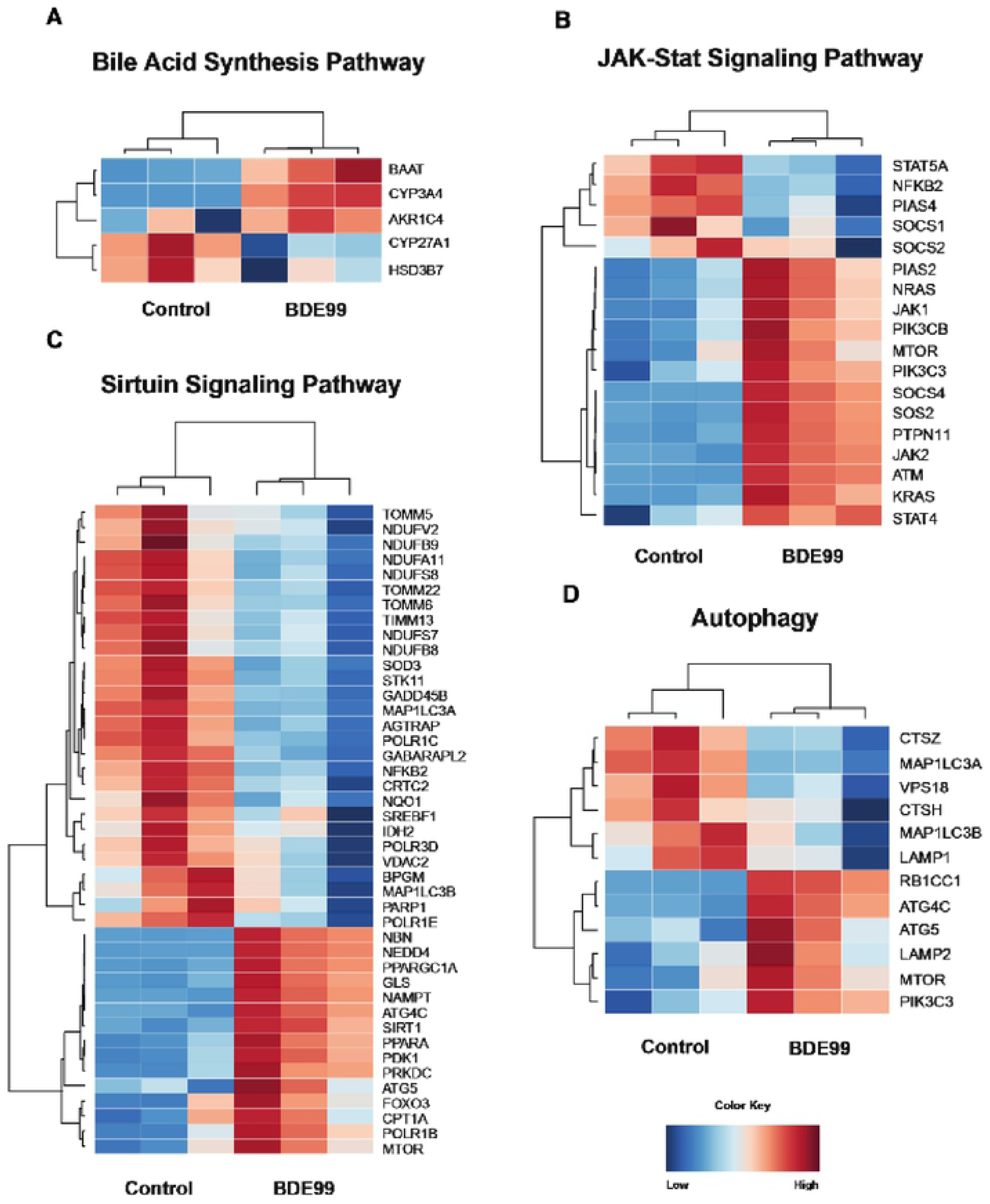
Effect of BDE-99: (A-D) Heatmaps depicting gene expression patterns of differentially expressed genes from BDE-99 exposure associated with bile acid synthesis, JAK-Stat signaling, sirtuin signaling, and autophagy. Blue indicates down­ regulation while red indicates up-regulation.

#### JAK-Stat Signaling Pathway

Overall, exposure to BDE-99 led to an up-regulation of genes involved in the JAK-Stat Signaling Pathway Figure (Figure 8B). Of the 18 differentially regulated genes, 13 were up-regulated whereas 5 were down-regulated. Suppressor of cytokine signaling 1 (SOCS1) and suppressor of cytokine signaling 2 (SOCS2), both members of the STAT-induced STAT inhibitor family, were down-regulated, while Janus kinase 1 (JAK1) and Janus kinase 2 (JAK2), tyrosine kinases that are involved in cell growth and cell development and are inhibited by SOCS1 and SOCS2, were up-regulated. Surprisingly, expression of Suppressor of cytokine signaling 4 (SOCS4), was up-regulated. Interestingly, Signal transducer and activator of transcription 4 (STAT4), part of the STAT family of transcription factors, was up-regulated while PIAS4 (Protein Inhibitor of Activated STAT4) was down-regulated.

Many genes encoding ras proteins such as SOS Ras/Rho Guanine Nucleotide Exchange Factor 2 (SOS2), K-ras proto-oncogene GTPase (KRAS), and N-ras proto-oncogene GTPase (NRAS) were up-regulated following exposure to BDE-99. The ras subfamily is involved in transmitting signals within cells and belongs to the small GTPase class of proteins. Other prominent genes were also up-regulated: PIK3CB and PIK3C3 (catalytic subunits of the PI3K complex), ATM serine/threonine kinase (master controller of cell cycle checkpoint signaling pathways), and mechanistic target of rapamycin (mTOR, central regulator of cellular metabolism).

#### Sirtuin Signaling Pathway and Autophagy

Sirtuins are a class of proteins that possess either mono-ADP-ribosyltransferase or deacylase activity. They regulate important cellular processes like transcription, apoptosis, aging, and inflammation in bacteria, archaea and eukaryotes. BDE-99 appears to down-regulate genes involved in oxidative phosphorylation (Figure 8C). In particular, many genes encoding subunits for Type I NADH dehydrogenase (NDUFV2, NDUFB9, NDUFA11, NDUFS8, NDUFS7, NDUFB8) were down-regulated in BDE-99 exposed cells. This enzyme is responsible for catalyzing the transfer of electrons from NADH to coenzyme Q10 and the first enzyme of the mitochondrial electron transport chain (Brandt, 2006). Furthermore, genes encoding proteins in the TIM/TOM complex (TOMM5, TOMM22, TOMM6, TIMM13) were down-regulated. This protein complex translocates proteins involved in oxidative phosphorylation through the mitochondrial membrane (Hood et al., 2003). In addition, genes involved in transcription, such as RNA polymerase I subunit B (POLR1B), and Protein kinase, DNA-activated, catalytic polypeptide (PRKDC), were up-regulated by treatment of BDE-99.

Several genes associated with autophagy and apoptosis were up-regulated and there was a large overlap in genes associated with autophagy and the sirtuin signaling pathway (Figure 7C and 7D). Neural precursor cell expressed developmentally down-regulated protein 4 (NEDD4) is a HECT ubiquitin ligase that accepts ubiquitin from an E2 ubiquitin-conjugating enzyme and transfers it to the target substrate, which is then marked for degradation. ATG4C, or autophagy related 4C cysteine peptidase, destroys endogenous proteins and damaged organelles. Although not essential in normal conditions, ATG4C is required for autophagic response during stressful conditions (Korkmaz et al., 2012). ATG5 is responsible for autophagic vesicle formation through an ubiquitin-like conjugating system (Yousefi et al., 2006). Forkhead box O3 (FOXO3) functions as a signal for apoptosis and transcribes genes necessary for cell death (Skurk et al., 2004).

There was a down-regulation of PCGs associated with tumorigenesis such as cathepsin Z (CTSZ), lysosomal associated membrane protein 1 (LAMP1), and cathepsin H (CTSH). CTSZ is a lysosomal cysteine proteinase and is often expressed ubiquitously in cancer cell lines (Jechorek et al., 2014). Increased expression of CTSH, in particular, has been found in the malignant progression of prostate cancer (Jevnikar et al., 2013). LAMP1 encodes a glycoprotein and is responsible for maintaining lysosomal integrity. The expression of LAMP1 has been observed on the surface of tumor cells from highly metastatic cancers, suggesting its role in the progression of lung cancer, colon cancer, and melanoma (Agarwal et al., 2015). Surprisingly, LAMP2, another glycoprotein closely related to LAMP1, was up-regulated in BDE-99 exposed cells (Figure 8D).

### Enriched canonical pathways following exposure to BDE-47, BDE-99, and CITCO

Since there is a large overlap in genes between BDE-47, BDE-99, and CITCO (Figure 2), it is suggested that these compounds may influence similar canonical pathways.

#### Interferon Signaling

Interferons are a group of signaling proteins that are produced and released by host cells in response to the presence of certain pathogens like bacteria, virus, and even tumor cells. There was a down-regulation in genes involved in apoptosis and antiviral properties such as IRF1, IRF9, BAK1, and MX1 in BDE-47, BDE-99, and CITCO groups (Figure 9A). Interferon regulatory factor 1 and 9 (IRF1 and IRF9) are transcriptional regulators of interferon-B and tumor suppressors of IFN-inducible genes. These factors modulate cell growth and help develop T cell immune responses to cancer cells (Harada et al., 1998). BCL2 antagonist/killer 1 (BAK1) plays a role in the mitochondrial apoptotic process, promoting the mitochondrial outer membrane permeability and releases apoptogenic factors into the cytosol (Chittenden et al., 1995). Interferon-induced GTP-binding protein (MX1) inhibits multiplication of the influenza virus and places the cell in a specific antiviral state (Chittenden et al., 1995).

**Figure 9:**
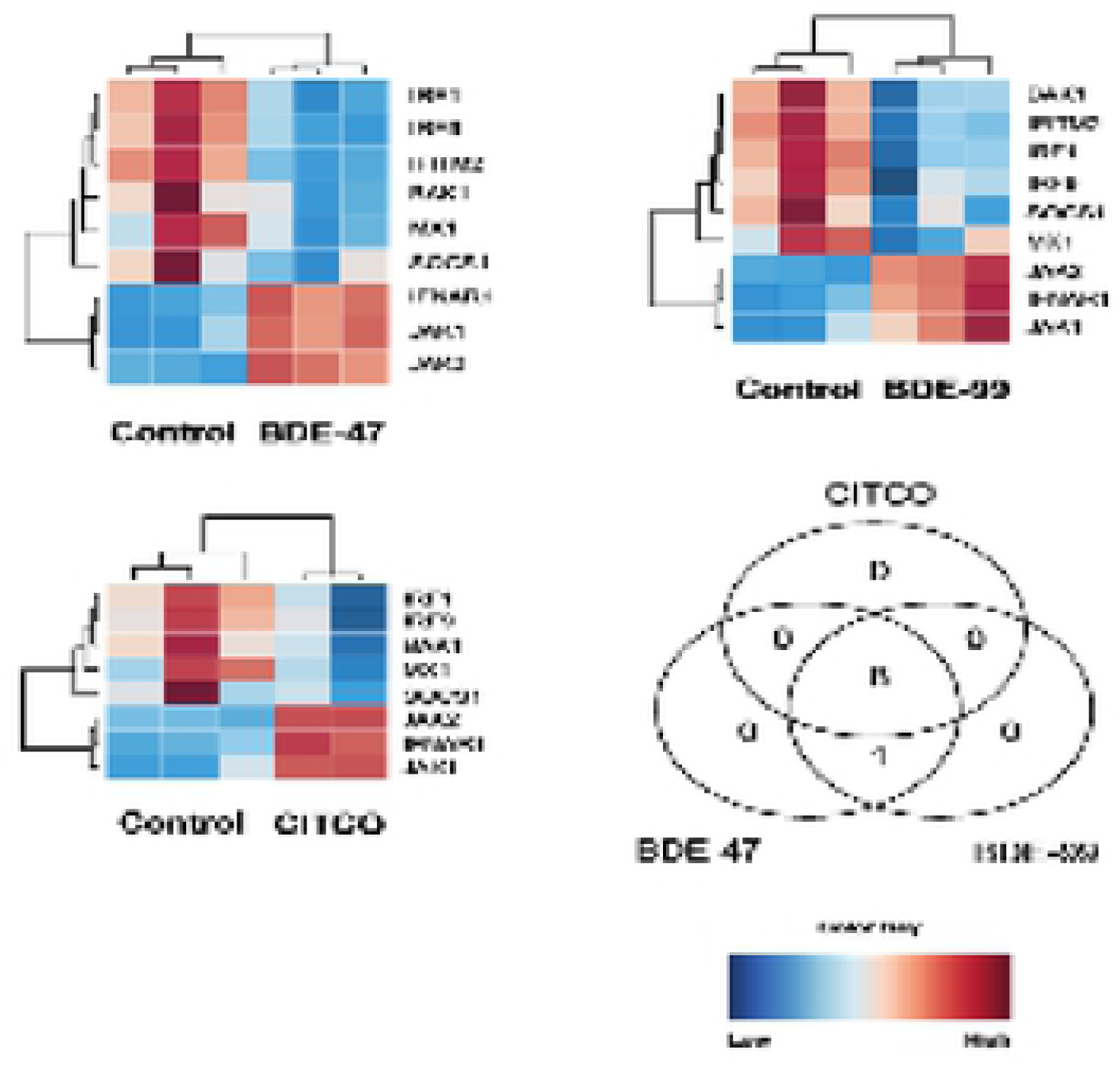
Effect of PBDEs and CITCO on interferon signaling: (A-C) Heat maps depicting gene expression of differentially expressed lncRNA-PCG pairs involved in interferon signaling. (D) Three-way Venn Diagram displaying the common and unique genes differentially expressed by CITGO, BDE-47, and BDE99 for interferon signaling. Blue indicates down-regulation while red indicates up-regulation

There was an up-regulation in genes involved in Jak-STAT signaling (*see Jak-STAT signaling*) following exposure to BDE-47, BDE-99, and CITCO. Janus kinase 1 and Janus kinase 2 (JAK1 and JAK2) were up-regulated while STAT-induced STAT inhibitor, suppressor of cytokine signaling (SOCS1) was down-regulated. Furthermore, interferon alpha and beta receptor subunit 1 (IFNAR1), a gene involved in the binding and activation of receptors that stimulate Janus protein kinases, was up-regulated as well (Figure 9A).

#### Glutathione Biosynthesis

Glutathione is an important antioxidant and protects against toxic xenobiotics involving oxidative stress and free radical intermediates. It is considered one of the most important molecules that cells can use to detoxify drugs and other toxins since it is both a nucleophile and a reductant (Lu, 2013). There was an up-regulation in glutathione biosynthesis in BDE-47, BDE-99, and CITCO groups (Figure 10A-B). The expression of glutamate-cysteine ligase regulatory subunit (GCLM) and glutamate-cysteine ligase catalytic subunit (GCLC) was up-regulated compared to the control. Both GCLM and GCLC are involved in the first rate-limiting step of glutathione biosynthesis and form the glutamate cysteine ligase. Glutamate cysteine ligase takes cysteine and glutamate to form γ-glutamylcysteine (Gipp et al., 1992).

**Figure 10:**
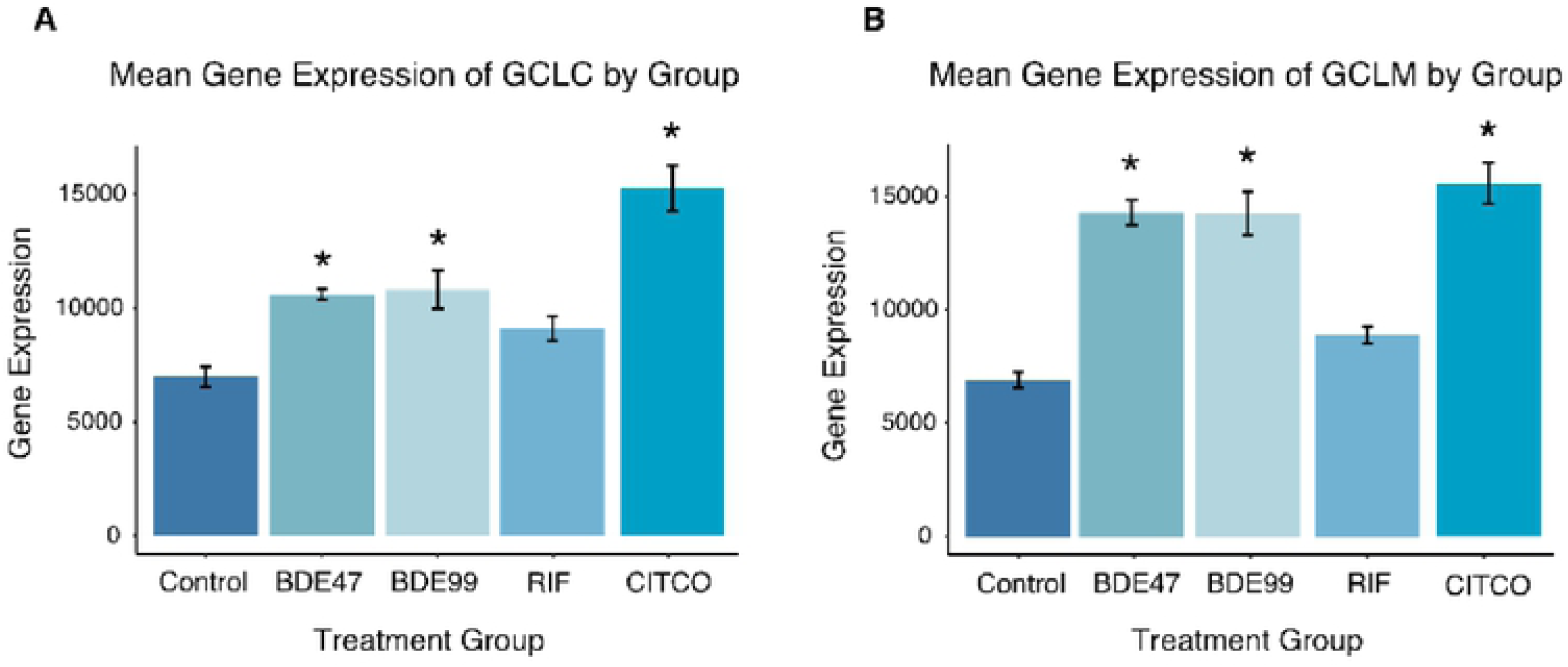
Effect of PBDEs and CITCO on glutathione biosynthesls: (A) Expression of GCLC by exposure group (B) Expression of GCLM by exposure group

#### mTOR signaling

The mechanistic target of rapamycin (mTOR) is a master growth regulator and plays an essential role in cellular processes, such as cytoskeletal organization, protein synthesis, and ribosomal biogenesis. mTOR is a critical component in two distinct protein complexes: mTOR complex 1 (mTORC1) and mTOR complex 2 (mTORC2). mTORC1 is activated by extra- and intracellular cues and stimulates cell growth and proliferation whereas mTORC2 phosphorylates enzymes important for cell survival and cytoskeletal organization (Dowling et al., 2010).

A number of ribosomal proteins (RPS3A, RPS27, RPS4X, RPS13, RPS9, RPS10, RPS29, RPS6, RPS15, RPS15A) and eukaryotic translation initiation factors (EIF4EBP1, EP3B, EIF3F, EIF4A3, EIF3L) were down-regulated after treatment with BDE-47, BDE-99, and CITCO, suggesting a decrease in translation capacity (Figure 11A). Although a majority of the ribosomal proteins were down-regulated, RPS6KA3, RPS6KB1, and RPS6KC1 were up-regulated. In addition to ribosomal proteins, phospholipase D2 (PLD2) was down-regulated; PLD2 catalyzes the hydrolysis of phosphatidylcholine to phosphatidic acid and choline and may be involved in cytoskeletal organization and transcriptional regulation (Ghim et al., 2016). The expression of serine/threonine kinase 11 (STK11) and protein kinase AMP-activated non-catalytic, proteins that both maintain cell metabolism, were down-regulated as well (Brown et al., 2011; Cheung et al., 2000).

**Figure 11:**
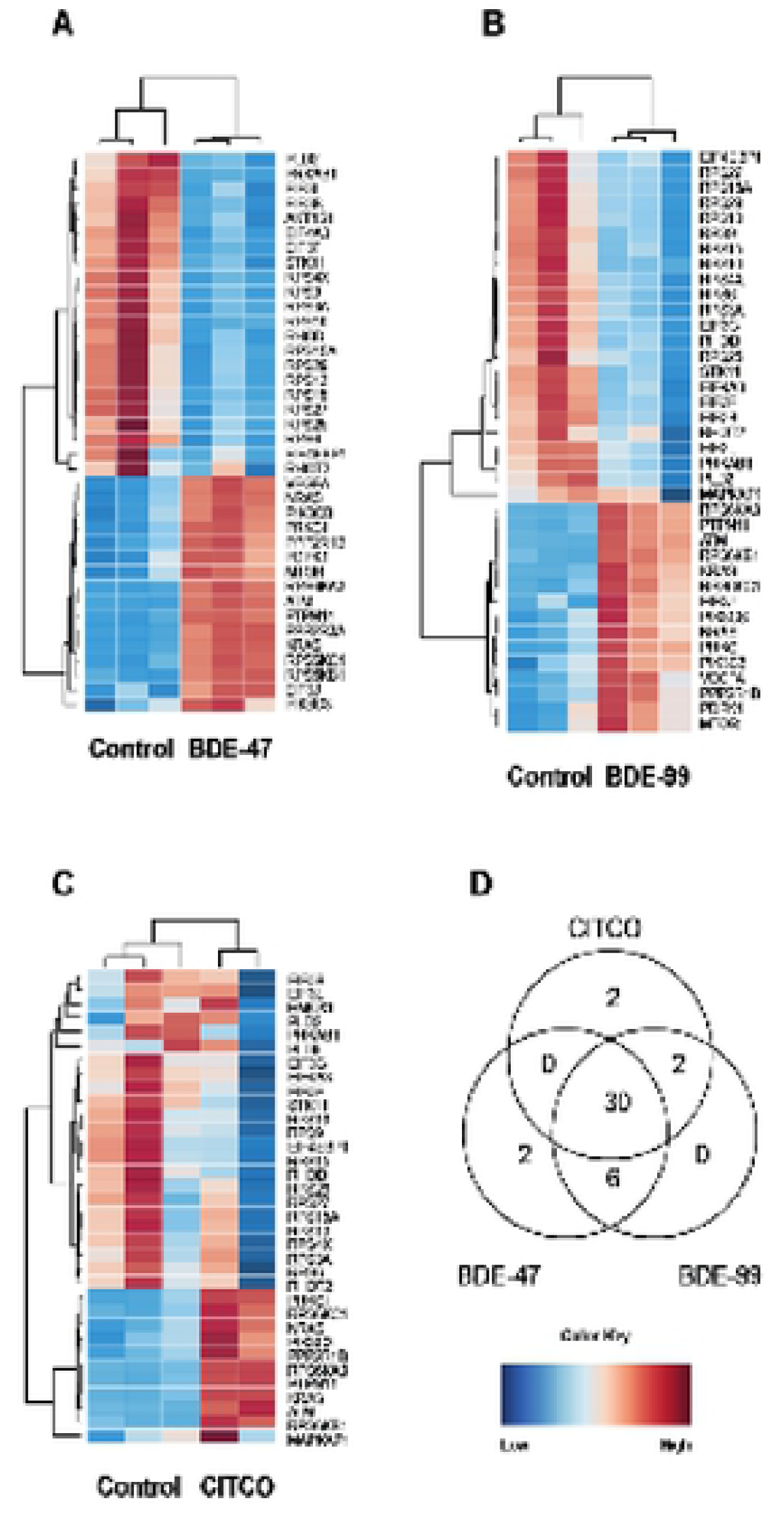
Effect of PBDEs and CITCO on mTORsignaling: (A-C) Heat maps depicting gene expression of differentially expressed lncRNA-PCG pairs involved in mTOR signaling. (D) Three-way Venn diagram showing overlapping and unique genes between the three groups. Blue indicates down­ regulation while red indicates up-regulation.

A number of genes involved in the Jak-STAT signaling pathway (KRAS, PIK3C3, ATM, NRAS, PTPN11, see *Jak-STAT Signaling*) were up-regulated following treatment to CITCO and the PBDEs (Figure 11A). Protein kinase C Iota (PRKCI) and protein phosphatase 2 scaffold subunit Abeta (PPP2R1B) were up-regulated as well. PRKCI is a member of the protein kinase C (PRK) family of serine threonine protein kinases and is shown to be a possible proto-oncogene (Qu et al., 2016). PPP2R1B is a constant regulatory subunit of protein phosphatase 2 and is a known tumor suppressor gene (Chou et al., 2007).

There were several genes that were expressed uniquely in BDE-47 and CITCO. AKT1 substrate 1 (AKT1S), a subunit of mTORC1, was down-regulated while protein phosphatase 2 regulatory subunit 3 Alpha (PPP2R3A) was up-regulated in BDE-47 (Figure 11B). The dysregulation of genes caused by BDE-47 exposure is comparable to the pattern caused by other chemical exposures: there is a decrease in mTOR signaling but an increase in expression of regulatory units that suppress the pathway. Phospholipase D family member 6 (PLD6) and heme oxygenase 1 (HMOX1) were down-regulated in CITCO-treated groups (Figure 11D). Since PLD2 was down-regulated in all three groups, it is not unusual for a related gene from the same superfamily, PLD6, to be down-regulated as well. Interestingly, HMOX1 is a gene essential in heme catabolism and cleaves heme to form biliverdin (Kwon et al., 2015).

### Validation of LncRNA-PCG pairs through LncTar

Initially, lncRNA-PCG pairs were determined if they were proximal to each other. LncTar, a tool that takes advantage of free-energy associations between RNA transcripts, was used on 25 drug-processing genes and their respective lncRNA pair. There were 50 lncRNAs that paired with 20 PCGs. Out of this initial list, 9 PCGs and 19 lncRNAs were validated using LncTar with a cutoff of −0.08 ndG. Multidrug resistance protein 3 (ABCB4) was paired with NONHSAT254289.1 and NONHSAT254288.1. ABCB4 is a member of the ATP-binding cassette (ABC) transporters and helps move phospholipids across the membranes of liver cells (Davit-Spraul et al., 2010). Another ABC transporter, multidrug resistance-associated protein 2 (ABCC2) was paired with NONHSAT158443.1 and NONHSAT158443.1. ABCC2 transports phase II products of biotransformations, i.e. conjugates of lipophilic substances with glutathione, glucuronate, and sulfate, in order to detoxify and protect the body (Jedlitschky et al., 2006). Aryl hydrocarbon receptor (AHR) paired with NONHSAT213218.1, NONHSAT213218.1, NONHSAT213218.1, and NONHSAT213218.1. AHR has been shown to regulate xenobiotic-metabolizing enzymes such as CYP1A1 (Barouki et al., 2007). N-acetyltransferease 8 (NAT8) paired with NONHSAT242202.1. NAT8 catalyzes the transfer of acetyl groups from acetyl-CoA to arylamines. Acetyl groups are important in the conjugation of metabolites from the liver and are essential to the metabolism and excretion of drug product (Veiga-da-Cunha et al., 2010). Nuclear receptor coactivator 7 (NCOA7), an estrogen receptor associated protein with punitive proteins in oxidation resistance, paired with NONHSAT208003.1 (Durand et al., 2007).

Several PCGs discussed earlier paired with lncRNAs according to LncTar. Genes involved in the PPARα/RXRα pathway (see *PPARα/RXRα pathway*) such as NCOR1 paired with NONHSAT145885.1; PPARA paired with NONHSAT245297.1, NONHSAT245299.1, NONHSAT245298.1, NONHSAT245301.1 and NONHSAT245300.1. GCLM (see *Glutathione biosynthesis*) paired with NONHSAT151954.1. Sirtuin 1 (SIRT1, see *Sirtuin signaling pathway*) paired with NONHSAT155619.1 and NONHSAT155619.1.

## DISCUSSION

In this study, we have presented evidence that PBDEs regulate both PCGs and lncRNAs in HepaRG cells in both PXR/CAR-ligand-dependent and independent manners. The majority of the differentially expressed lncRNAs are mapped to the introns of PCGs, indicating that they may be produced from post-transcriptional splicing of nascent mRNA transcripts. Furthermore, PBDEs regulate PCG-lncRNA pairs involved in protein ubiquitination, PPARα-RXRα activation, carbohydrate metabolism, mTOR signaling, glutathione biosynthesis and interferon signaling. BDE99, specifically, modulates PCG-lncRNA pairs involved in bile acid synthesis, cell survival and death as well as autophagy. In addition, we validated distinct PCG-lncRNA pairs involved in xenobiotic biotransformation through lncTar.

We identified a total of 13,204 genes that were uniquely expressed by at least one of the four exposures groups. There were 8367 PCGs shared between the PBDEs and CITCO; whereas there were fewer PCGs (3825) shared between the PBDEs and RIF. With regards to lncRNAs, CITCO and PBDEs shared 4610 genes while RIF and the PBDEs shared 1812 genes. Consistent with the result in this study, previous studies have shown that PBDEs activate PXR/CAR pathways in mice (Li et al., 2018). With regards to CAR pathway mediation, a previous study demonstrated the activation of CAR pathways through BDE-47 in human primary hepatocytes (Sueyoshi et al., 2014). Our result is novel: there has been no literature yet showing that CAR activation by PBDES is more prominent than PXR activation.

Through histone methylation and the subsequent change in chromatin structures, lncRNAs can act as epigenetic modifiers of transcription. The lncRNA HOTAIR contributes to the demethylation of H3 lysine-4 (H3K4), leading to the suppression of transcriptional activation. Urothlial cancer associated 1 (UCA1) is a lncRNA that acts as a post-transcriptional inhibitor of cell cycle regulation. UCA1 competitively binds to heterogenous ribonucleoprotein I (hnRNP 1), which is required for the translation of the tumor suppressor gene p27 (Dempsey and Cui, 2017). The circular lncRNA, ci-ankrd52, is a positive regulator of Pol II transcription, suggesting a cis-regulatory role of noncoding intronic transcripts on their parent coding genes (Zhang et al., 2013a). Although UCA1 and ci-ankrd52 were not regulated by PBDEs, they are intronic lncRNAs that have documented roles in transcription and may provide possible mechanisms for PBDE-regulated lncRNAs.

The expression of genes involved in protein ubiquitination were significantly altered in all four exposure groups. A previous study provided evidence that RIF increased the ubiquitin-proteasome degradation of MRP2 in HepG2 cells through the E3 ubiquitin ligase GP78. Furthermore, we demonstrated that oxidative stress induced by RIF activated the ERK/JNK/p38 and PI3K signaling pathways, thus resulting in clathrin-dependent endocytosis (Xu et al., 2019). With regards to the relationship between protein ubiquitination and CITCO, previous literature has shown that even after CITCO exposure, proteasomal inhibition disrupted CAR function by repressing CAR nuclear trafficking and inhibiting induction of CAR target gene responses in human primary hepatocytes (Chen et al., 2014). Since the PBDE exposed groups had 15 genes and 28 genes in common with the RIF and CITCO treatment group, respectively, BDE-47 and BDE-99 may affect protein ubiquitination pathways through similar mechanisms modulated by RIF and CITCO.

The PPARα/RXRα pathway was also significantly altered in the BDE-47, BDE-99, RIF, and CITCO exposed groups. The PPARα/RXRα heterodimer is necessary for the activation of PPARα, a major transcription factor involved in lipid metabolism. PBDEs restrict fatty acid esterification by suppressing the activity of a key enzyme, phosphoenolpyruvate carboxykinase (PEPCK). The suppression of PEPCK activity resulted in altered lipid metabolism through the redirection of fatty acids away from esterification and towards the synthesis of ketones (Cowens et al., 2015). In this study, however, many genes involved in the PPARα/RXRα pathway, specifically in cell bioenergetics, were up-regulated by all four exposure groups, suggesting the promotion of ketogenesis. In a separate study, exposure to BDE-47 up-regulated the expression of cluster of differentiation 36 (CD36) in CD-1 mice. Increased expression of Cd36 caused an increase in triglyceride levels, suggesting that Cd36 plays a role in the blood/liver imbalance of triglycerides. In our current study, Cd36 was decreased in HepaRG cells exposed to BDE-47 and concurs with our hypothesis that PBDEs may increase ketones through the metabolism of triglycerides (Khalil et al., 2018). In mice fed a high-fat diet, however, BDE-47 caused an increase of body weight and inflammation through the increased synthesis of triglycerides. In addition to obesity, exposure to BDE-47 may lead to detrimental hepatotoxicity through hepatic steatosis and liver fibrosis (Yang et al., 2019).

Genes involved in GDP-L-fucose biosynthesis I were down-regulated in the BDE-47 and BDE-99 treatment groups but not in the RIF or CITCO treatment groups, suggesting that PBDEs play a role in carbohydrate metabolism unique from PXR/CAR pathways. Our finding of intermediary metabolism distruption coincides with previous studies; a paper in 2019 demonstrated that mice exposed to BDE-47 and BDE-99 had significant alterations in metabolic syndrome-related intermediary metabolites in serum, the liver and small intestinal contents (Scoville et al., 2019). Among PBDE-regulated metabolites analyzed in the paper, lower levels of mannose were reported in mice exposed to BDE-99. Since we observed the down-regulation of genes involved in converting GDP-D-mannose to GDP-L-fucose, we would expect higher levels of mannose in a metabolite sample.

Bile acid synthesis, sirtuin signaling pathway, and autophagy were uniquely altered by exposure to BDE-99. A 2018 study demonstrated that BDE-99 increased many unconjugated bile acids in the serum, liver, small intestinal content, and large intestinal content of mice with conventional microbiomes but not in germ-free mice, suggesting a microbiome-dependent effect of BDE-99. These increasing levels of bile acids corresponded to an increased abundance of bacterial and their resulting microbial enzymes. Unconjugated bile acids are generally thought to be more toxic than their conjugated counterparts; an excess of unconjugated bile acids could lead to impaired liver function and cholestasis (Li et al., 2018). With regards to the sirtuin signaling pathway, exposure to pollutants can affect SIRT1 expression; this can affect the expression of downstream proteins and result in toxic damage. Upregulation of SIRT1 expression is protective against the toxicity of pollutants ((Li et al., 2018). In our current study, SIRT1 was significantly down-regulated in HepaRG cells exposed to BDE-99, suggesting that this PBDE can leave hepatocytes vulnerable to toxic attacks.

The results from our PCG-lncRNA study support previous work conducted in mice exposed to BDE-47 and BDE-99. Specifically, in our study, lncTar predicted that the binding of lncRNAs with ABCB4, ABCC2, AHR, GCLM, NAT8, NCOA7, NCOR1, PPARA, and SIRT1 (Table 2). LncRNAs paired with ABCC2 and NCOR1 were differentially regulated in the livers of germ-free mice (Li et al., 2018). In addition to supporting our findings, these results suggest that certain lncRNA-PCG interactions are possibly conserved between the two species.∂

From the gene pathway analysis, we speculate that PBDEs may exert their hepatotoxicity through mitochondrial dysfunction and inflammation. For example, several TOMM/TIMM genes were significantly down-regulated by BDE-99 (Figure 10). PBDEs can cause mitochondrial dysfunction by inhibiting the electron transport chain, altering mitochondrial morphology, increasing apoptosis, and increasing oxidative stress; several of these events have been reported in individuals with autism spectrum disorder (Li et al., 2018). Although there is little information concerning the effect of BDE-99 on autophagy, an increase in autophagic activity has been documented in HepG2 cells to following exposure to BDE-100. Following BDE-100 exposure, HepG2 cells had increased staining with lysosomal dye. The study also noted that the mitochondrial DNA copy number decreased in these cells, signifying an attempt from the cell to manage mitochondrial damage by selective mitophagy (Pereira et al., 2017).

In addition, inflammation-related signaling pathways involving interferons, mTOR and Jak-STAT were altered after PBDE exposure. There was a positive relationship between PBDEs and IL-6 and TNF in a study assessing the relationship between PBDEs and inflammation biomarkers in pregnancy and post-partum (Zota et al., 2018). Exposure to BDE-47 was shown to cause inflammatory cell infiltration in the liver and increased levels of a Kupffer cell marker and proinflammatory cytokines and chemokines. Additionally, BDE-47 exposure significantly increased acetylation of p65 and H3K9 which resulted in the subsequent transcription of inflammation-related gens in the livers of mice (Zhang et al., 2015). While the expression of mTOR was up-regulated in this study, many ribosomal proteins were down-regulated; this may be a compensatory action in preventing the activation of inflammation pathways following exposure to toxicants.

PBDEs are known to cause oxidative stress and the effects of BDE-47 have shown to be more pronounced in GCLM(-/-) cultured cells than in GCLM(+/+) cells, suggesting that the synthesis of glutathione protects the cell from PBDE-induced toxicity by increasing antioxidant activity. (Fernie et al., 2005). Since we observed increased expression of genes involved in GSH synthesis following exposure to BDE-47, BDE-99, and CITCO, we hypothesize that this may be a compensatory response. A mouse study published in 2016 showed that the expression of GCLC, which is involved in the synthesis of GSH, was increased through CAR activation but not PXR activation (Cui and Klaassen, 2016). The expression of GCLC was also increased in BDE-47 and BDE-99, suggesting that PBDEs affect glutathione synthesis through CAR mediated pathways. Since PBDEs induce hepatic oxidative stress, it has been shown that PBDE exposure increases the glutathione disulfide/GSH ratio and the levels of oxidized glutathione in the liver (Fernie et al., 2005). An increase in glutathione synthesis suggests a protective effect against toxicant exposure.

There were several limitations of this study. LncRNAs were detected through poly A tail selection, leading to possible bias. The poly A tail is not a universal trait of all lncRNAs; as a result, we were not able to quantify all the lncRNAs in our sample. This is evidenced in Figure 2, where a majority of lncRNAs were not detected. Follow-up studies should be conducted to verify the differentially-regulated pathways by PBDE exposure.

In conclusion, this study is the first to characterize the relationship between lncRNAs and PCGs after exposure to PBDEs in HepaRG cells. We demonstrated that the expression of number of lncRNAs are significantly altered by BDE-47 and BDE-99 in both PXR/CAR-ligand-dependent and independent manners. Further research into the roles of these RNA molecules will be vital for the understanding the overall genomic changes enacted by the toxicant exposure.

## ACKNOWLEDGMENTS

Supported by UW BEBTEH Training Program, NIH R01 grants GM111381, ES025708, University of Washington Center for Exposures, Diseases, Genomics, and Environment [Grant P30 ES0007033], and the Murphy Endowment).

